# Quantitative Seq-LGS – Genome-Wide Identification of Genetic Drivers of Multiple Phenotypes in Malaria Parasites

**DOI:** 10.1101/078451

**Authors:** Hussein M. Abkallo, Axel Martinelli, Megumi Inoue, Abhinay Ramaprasad, Phonepadith Xangsayarath, Jesse Gitaka, Jianxia Tang, Kazuhide Yahata, Augustin Zoungrana, Hayato Mitaka, Paul Hunt, Richard Carter, Osamu Kaneko, Ville Mustonen, Christopher J. R. Illingworth, Arnab Pain, Richard Culleton

## Abstract

Identifying the genetic determinants of phenotypes that impact on disease severity is of fundamental importance for the design of new interventions against malaria. Traditionally, such discovery has relied on labor-intensive approaches that require significant investments of time and resources. By combining Linkage Group Selection (LGS), quantitative whole genome population sequencing and a novel mathematical modeling approach (qSeq-LGS), we simultaneously identified multiple genes underlying two distinct phenotypes, identifying novel alleles for growth rate and strain specific immunity (SSI), while removing the need for traditionally required steps such as cloning, individual progeny phenotyping and marker generation. The detection of novel variants, verified by experimental phenotyping methods, demonstrates the remarkable potential of this approach for the identification of genes controlling selectable phenotypes in malaria and other apicomplexan parasites for which experimental genetic crosses are amenable.

**Significance Statement:** This paper describes a powerful and rapid approach to the discovery of genes underlying medically important phenotypes in malaria parasites. This is crucial for the design of new drug and vaccine interventions. The approach bypasses the most time-consuming steps required by traditional genetic linkage studies and combines Mendelian genetics, quantitative deep sequencing technologies, genome analysis and mathematical modeling. We demonstrate that the approach can simultaneously identify multigenic drivers of multiple phenotypes, thus allowing complex genotyping studies to be conducted concomitantly. This methodology will be particularly useful for discovering the genetic basis of medically important phenotypes such as drug resistance and virulence in malaria and other apicomplexan parasites, as well as potentially in any organism undergoing sexual recombination.

## INTRODUCTION

Malaria parasite strains are genotypically polymorphic, leading to a diversity of phenotypic characteristics that impact on the severity of the disease they cause. Discovering the genetic bases for such phenotypic traits can inform the design of new drugs and vaccines.

Experimental studies of the genetic basis of particular phenotypes have previously relied on linkage or association mapping. Linkage mapping studies involve the crossing of two inbred (or clonal) parental stains to produce one or more recombinant progenies, from which individual recombinant offspring are isolated and typed for the phenotype under investigation, while in association mapping studies unrelated individuals are sampled and phenotyped (1). Both approaches require the genotyping of individual progeny through the use of genetic markers, and in the case of linkage mapping, this requires the cloning of progeny following the production of a genetic cross. Prior to the advent of high-throughput deep sequencing using second and third generation sequencing technologies, the identification of loci associated with particular phenotypes relied on the development of large numbers of molecular markers through approaches such as restriction fragment length polymorphism (RFLP) (2), amplified fragment length polymorphism (AFLP) (3) or microsatellite markers (4).

Both association mapping and linkage analyses approaches have been adopted to understand the genetic mechanisms behind various phenotypes in malaria (5–9) and with the application of whole genome sequencing (WGS), the resolution of these methodologies has been dramatically improved, allowing the discovery of selective sweeps as they arise in the field (10).

However, both approaches suffer from drawbacks when working with malaria parasites: linkage mapping requires the cloning of individual recombinant offspring, a process that is both laborious and time-consuming, although the recent combination of genetically engineered parasites and flow cytometry has shown great potential to address the issue (11).

Association studies, on the other hand, require the collection of a large number of individual parasites (usually in the thousands) from diverse geographical origins and over periods of several months or years to produce enough resolution for the detection of selective sweeps. This approach also requires a considerable amount of manpower and resources.

Linkage Group Selection (LGS), like linkage mapping, relies on the generation of genetic crosses, but bypasses the need for extracting and phenotyping individual recombinant clones. Instead, it relies on quantitative molecular markers to measure allele frequencies in the recombinant progeny and identify loci under selection (12, 13). It has been successfully applied in studying strain-specific immunity (SSI) (14, 15), drug resistance (12, 16) and growth rate (Pattaradilokrat et al. 2009) in malaria and SSI in *Eimeria tenella* (17). More recently, LGS also benefited from the advent of quantitative WGS, facilitating the generation of molecular markers (the SNPs identified during WGS) and improving the resolution and accuracy of the approach (18).

An approach similar to the above, applied to yeast, has also shown that complex trait phenotypes, such as heat tolerance, can be analyzed, exploiting the possibility of collecting time-resolved sequence data from such a system (19, 20). However, in an experimental malaria system, collection of data is not so straightforward. LGS also lacks a formal mathematical methodology to distinguish selective sweeps (or “selection valleys”) from false positive signals (e.g. sudden shifts in allele frequency within a chromosome due to clonal growth in the cross population, or simply random background noise) or to accurately define the boundaries of regions of the genome under significant selection.

In the present study, LGS has been applied to a genetic cross between two strains of the rodent malaria parasite *Plasmodium yoelii* that display two discernible phenotypes, namely a difference in growth rate (with strain 17X1.1pp growing faster than strain CU) and the ability to induce protective immunity against homologous challenges with the immunizing strain. A sophisticated mathematical model, built upon methodological developments in the analysis of genetic cross populations (20), was developed to analyze the data.

This modified LGS approach relies on the generation and independent selection of two independent crosses between the same parental clone parasites that differ in the phenotype under investigation. The progeny from both crosses pre- and post-selection are then subjected to high depth WGS, and SNP marker movement analyzed using best fitting modeling (termed quantitative-seq LGS, qSeq-LGS), as summarized in **Figure 1**.

**Figure 1.**
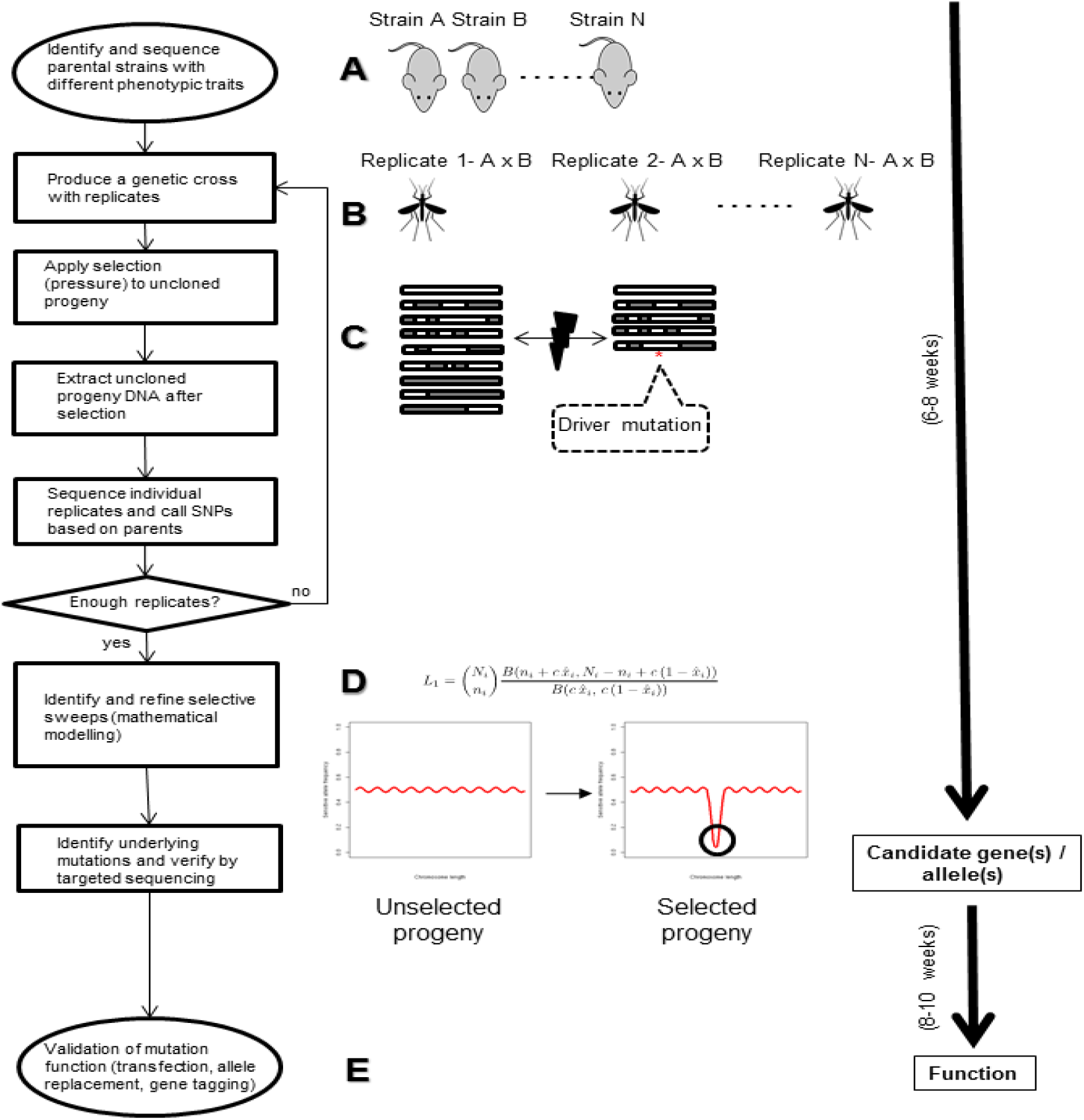
Graphical representation of the Quantitative-seq LGS approach. (**A**) The process starts with the identification of distinct selectable phenotypes in cloned strains of the pathogen population (in this case malaria parasites) and their sequencing, usually from the vertebrate blood stage. (**B**) Multiple genetic crosses between two cloned strains are produced, in this case inside the mosquito vector. (**C**) Each individual cross progeny is grown with and without selection pressure(s). Selection pressure will remove those progeny individuals carrying allele(s) associated with sensitivity to the selection pressure(s), while allowing progeny individuals with the resistant allele(s) to survive. DNA is then extracted from the whole, uncloned progeny for sequencing. SNPs distinguishing both parents are used to measure allele frequencies in the selected and unselected progenies. (**D**) A mathematical model is applied to identify and define selective sweeps. Selective sweep regions are then analyzed in detail to identify potential target mutations underlying the phenotype(s) being studied. Localized, capillary sequencing can be employed to verify or further characterize mutations. (**E**) Finally, where applicable, transfections studies, allele replacement experiments or other approaches can be carried out to confirm the effect of target mutations.

Applying this approach to crosses between two strains of *P. yoelii* that induce SSI, and which differ in their growth rates, we were able to identify three genomic regions containing genes controlling both phenotypes. Two genomic selection valleys were identified controlling SSI, and one controlling the growth rate differences. Of the SSI genomic valleys, one contained the gene encoding Merozoite Surface Protein 1 (MSP1) known to be a major antigen, and the other contained the *P. yoelii* orthologue of Pf34, a GPI-anchored rhoptry protein and potential antigen. Growth rate differences were found to be controlled by a newly described mutation in region 2 of the *P. yoelii* Erythrocyte Binding Ligand gene (*Pyebl*).

These results demonstrate that qSeq-LGS can be used to analyze multiple complex phenotypes concomitantly. We also introduce a mathematical model for the rapid discovery of selective sweeps within the genome and for the accurate definition of loci containing target genes.

The resulting method allows the identification of candidate alleles conferring a phenotype directly from the genomes within weeks and at high resolution, thus bypassing the limitations of more traditional genetic linkage studies.

## RESULTS

### Characterization of strain specific immunity and growth rate phenotypic differences between CU and 17X1.1pp

The difference in growth rate between the blood stages of the two clones was followed *in vivo* for nine days in CBA mice. A likelihood ratio test using general linear mixed models indicated a more pronounced growth rate for 17X1.1pp compared to CU (clone by time interaction term, L = 88.60, df = 21, P < 0.0001, **Figure 2A**). To verify that the two malaria clones could also be used to generate protective SSI, groups of mice were immunized with 17X1.1pp, CU or mock immunized, prior to challenge with a mixture of the two clones. The relative proportions of the two clones were measured on day four of the infection by real time quantitative PCR targeting the polymorphic *msp1* locus (21). A strong, statistically significant SSI was induced by both parasite strains in CBA mice (**Figure 2B**).

**Figure 2.**
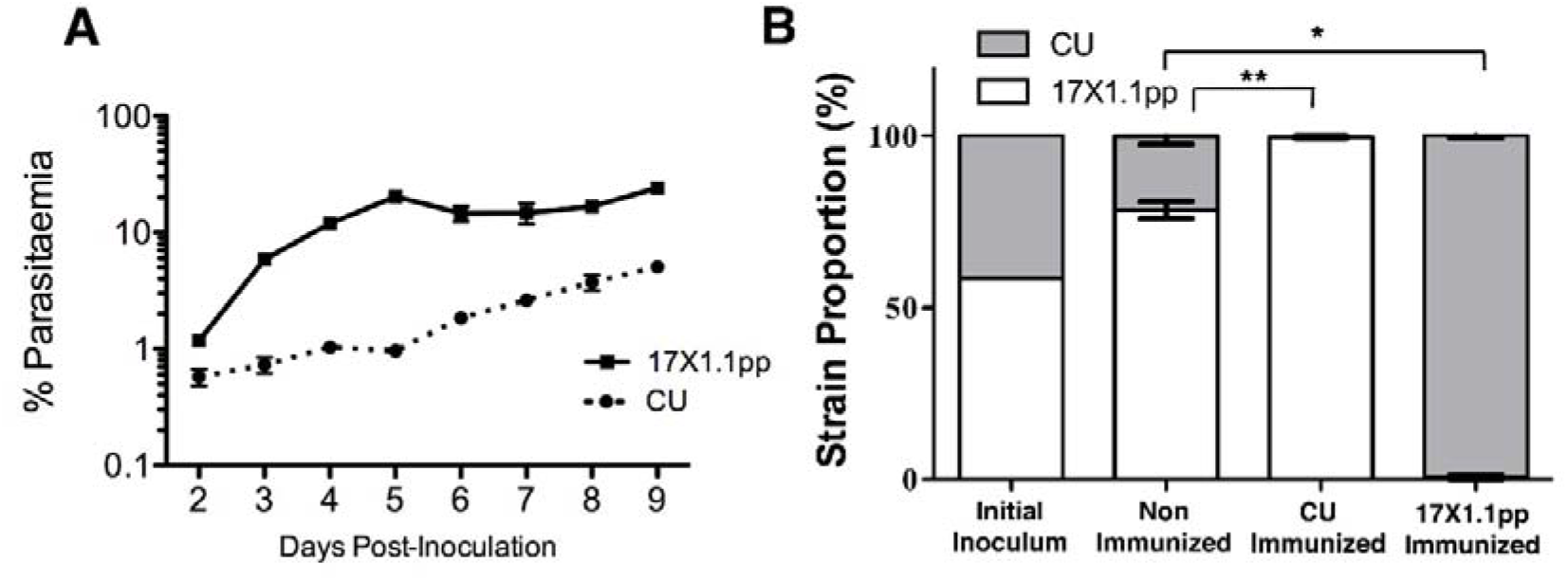
**(A)** Growth rate of *Plasmodium yoelii* strains 17X1.1pp and CU in CBA mice inoculated with 1 × 10^6^ iRBCs on Day 0. Error bars indicate the standard error of the mean for three mice per group. **(B)** The relative proportions of CU and 17X1.1pp were measured by RTQ-PCR targeting the polymorphic *msp1* locus at Day 4 post-inoculation with a mixed inoculum containing approximately equal proportions of both strains in naïve mice and mice immunized with one of the two strains. Error bars show the standard error of the mean of five mice per group. * p < 0.05, Wilcoxon rank sum test, W = 25, p = 0.0119, n = 5; ** p < 0.01, Wilcoxon rank sum test, W = 25, p = 0.0075, n = 5

### Identification of selection valleys

Two kinds of selection pressure were applied in this study: growth rate driven selection and SSI. Two independent genetic crosses between 17X1.1pp and CU were produced, and both these crosses subjected to immune selection (in which the progeny were grown in mice made immune to either of the two parental clones), and grown in non-immune mice. Progeny were harvested from mice four days after challenge, at which point strain specific immune selection in the immunized mice, and selection of faster growing parasites in the non-immune mice had occurred. Using deep sequencing by Illumina technology, a total of 29,053 high confidence genome-wide SNPs that distinguish the parental strains were produced by read mapping with custom-made Python scripts. These were filtered using a likelihood ratio test to remove sites where alleles had been erroneously mapped to the wrong genome location. Next a “jump-diffusion” analysis was applied to identify allele frequency changes (**Supplementary Table S1**) that were likely to have arisen from the clonal growth of individuals within the cross population or possible incorrect assembly of the reference genome, as described in the Methods section and in more detail in the Supplementary Protocol. Based upon an analytical evolutionary model describing patterns of allele frequencies following selection, a maximum likelihood approach was used to define confidence intervals for the positions of alleles under selection in each of the genetic cross populations (**Supplementary Tables S2-6)**.

A total of five potential selective sweeps were detected among all the samples (Table 1). Of these, two were detected in both replicates and across multiple progeny samples (**Figure 3A**). When considering the combined largest intervals, a selective sweep was located at position 1,436-1,529 kb on Chromosome (Chr) XIII and the other was located at position 1,229-1,364 kb on Chr VIII (**Figure 3B**).

**Figure 3.**
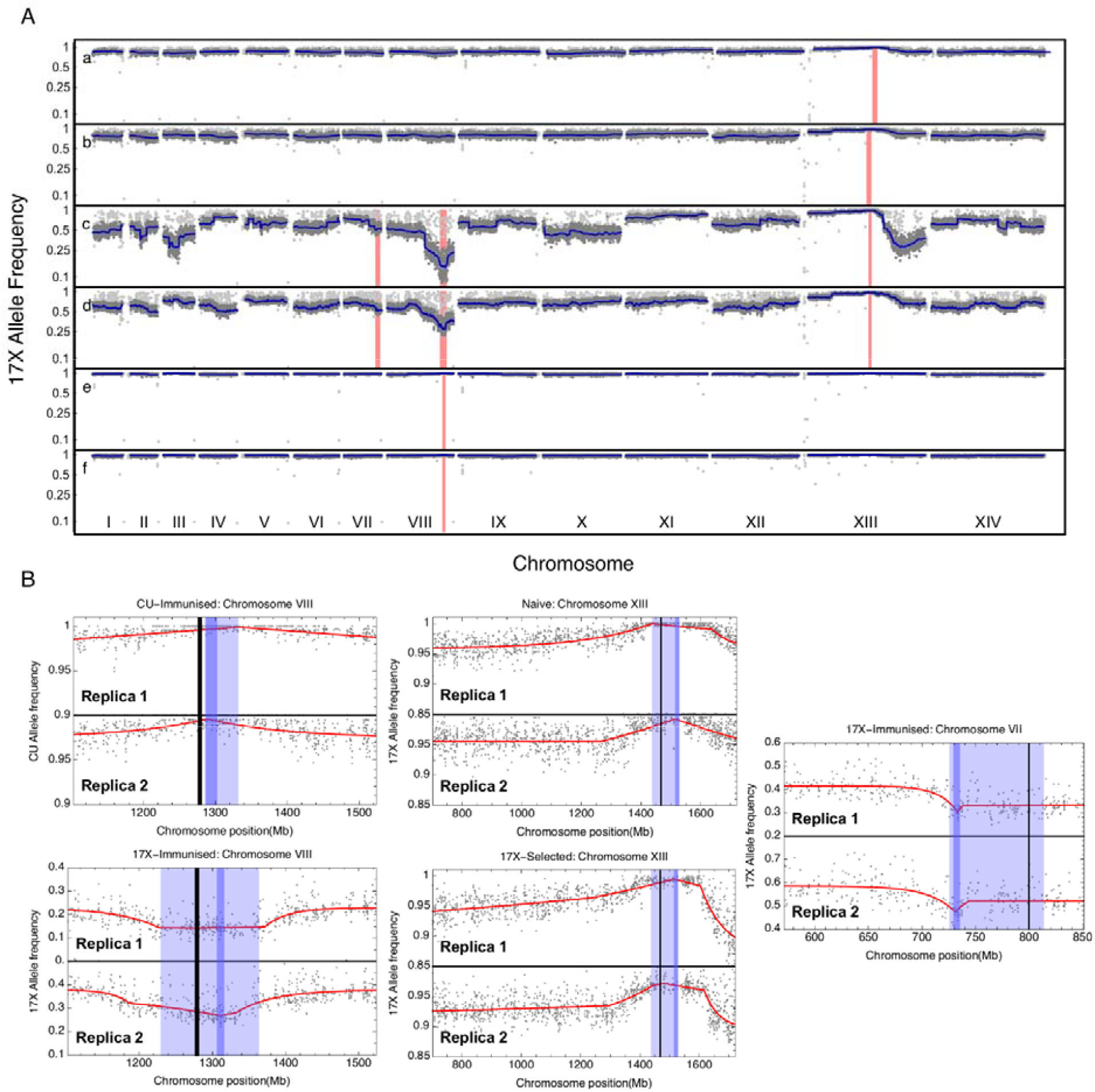
(**A**) Genome-wide *P. yoelii* CU allele frequency of two independent genetic crosses grown in (a,b) naïve mice, (c,d) 17X1.1pp immunized mice and (e,f) CU-immunized mice. Light gray dots represent observed allele frequencies. Dark gray dots represent allele frequencies retained after filtering. Dark blue lines represent a smoothed approximation of the underlying allele frequency; a region of uncertainty in this frequency, of size three standard deviations, is shown in light blue. A conservative confidence interval describing the position of an allele evolving under selection is shown via a red bar. (**B**) Evolutionary models fitted to allele frequency data. Filtered allele frequencies are shown as gray dots, while the model fit is shown as a red line. Dark blue and light blue vertical bars show combined and conservative confidence intervals for the location of the selected allele. A black vertical line shows the position of a gene of interest.

**Table 1.** Confidence intervals for driver locations as determined by mathematical modeling.

| **Condition** | **Selected Allele<sup>1</sup>** | **Allele location<sup>2</sup> (kb)** | **Combined interval (kb)<sup>3</sup>** | **Conservative interval (kb)<sup>4</sup>** |
| --- | --- | --- | --- | --- |
| Naïve | CU | XIII: 1513.899 | 1513.899 - 1528.275 | 1436.717 - 1528.275 |
| 17X1.1pp-selected | 17X1.1pp | VII: 733.215 | 729.368 - 735.525 | 725.528 - 813.866 |
| 17X1.1pp-selected | CU | VIII: 518.574 | 509.379 - 684.238 | n/a* |
| 17X1.1pp-selected | 17X1.1pp | VIII: 1311.291 | 1306.083 - 1316.428 | 1229.582 - 1363.920 |
| 17X1.1pp-selected | CU | XIII: 1480.699 | 1481.090 - 1489.964 | 1481.090 - 1528.275 |
| 17X1.1pp-selected | 17X1.1pp | XIII: 2256.431 | 2149.868 - 2264.801 | n/a* |
| CU-selected | CU | VIII: 1294.402 | 1286.595 - 1302.526 | 1285.184 - 1332.198 |

<sup>1</sup> indicates the direction of selection.
<sup>2</sup>The maximum likelihood location of the selected allele by chromosome genome coordinate.
<sup>3</sup>The combined interval was produced by combining the likelihoods calculated for each replica, with selected alleles having the same positions in each replica, ignoring biological noise.
<sup>4</sup>The conservative interval was defined as the minimal set of adjacent loci containing both of the replica-specific confidence intervals and takes biological noise into account. See Supplementary Methods for more details.
\* indicates a driver only found in a single replica

The selection valley on Chr XIII was the result of selection against CU-specific alleles at the target locus in both replicate crosses grown in non-immunized mice and replicate crosses grown in 17X1.1pp-immunised mice (**Figure 3A**).

The selective sweep on Chr VIII was detected exclusively in the parasite crosses grown in immunized mice. This selection valley was consistently detected in both replicates and in both CU and 17X1.1pp immunized mice. Furthermore, the selection pressure acted against both CU and 17X1.1pp alleles at corresponding loci (**Figure 3A** and **Supplementary Figure S1A**). The signal was weaker when measuring the allele frequency of the non-immunizing strain, but an effect was still observable (**Figure 3A** and **Supplementary Figure S1A**).

Of the remaining five selective sweeps, only one was detected across replicates in at least one of the samples, namely the locus between positions 725-814 kb on Chr VII (**Table 1 and Figure 3A and B**). This event was only detected in mice replicates immunized with the 17X1.1pp strain and significantly affected the proportion of 17X1.1pp alleles (**Supplementary Figure S1B**), although a corresponding increase in CU-allele frequencies was also apparent (**Figure 3A**).

The remaining selection valleys (on Chrs VIII and XIII) were only detected in a single sample and in a single replicate (**Table 1**) and were thus considered to be non-significant.

### Potential target genes within the three main selection valleys

The selection valley associated with SSI on Chr VIII contains 41 genes of which one, *msp1*, is a well characterized major antigen of malaria parasites that has been proposed as a vaccine candidate (22) and has been previously linked to SSI in *Plasmodium chabaudi* (14, 15). No other gene in this selection valley matched the profile of a potential antigen (based on functional description, presence of TM domains and/or signal peptides).

The growth rate associated selection valley on Chr XIII contains 29 genes, including *Pyebl*, a gene previously implicated in growth rate differences between strains of *P. yoelii* (24, 25).

Finally, the minor selection valley on Chr VII consists of 21 genes with several genes bearing signatures associated with a potential antigenic role.

### Characterization of EBL as the major driver of growth rate differences through alleleic replacement

Due to its location in the selection valley associated with growth rate selection and its previous identification as a gene underlying multiplication rate differences in *P. yoelii*, *Pyebl* was selected for further analysis. Sanger capillary sequencing re-confirmed the existence in 17X1.1pp of an amino acid substitution (Cys -> Tyr) at position 351 within region 2 of the gene. When aligned against other *P. yoelii* strains and other *Plasmodium* species, this cysteine residue is highly conserved, thus making the substitution that arose in 17X1.1pp unique (**Figure 4**). Crucially, no other mutations were detected in the coding sequence of the gene, including in region 6, the location of the SNP previously implicated in parasite virulence in other strains of *P. yoelii* (23).

**Figure 4.**
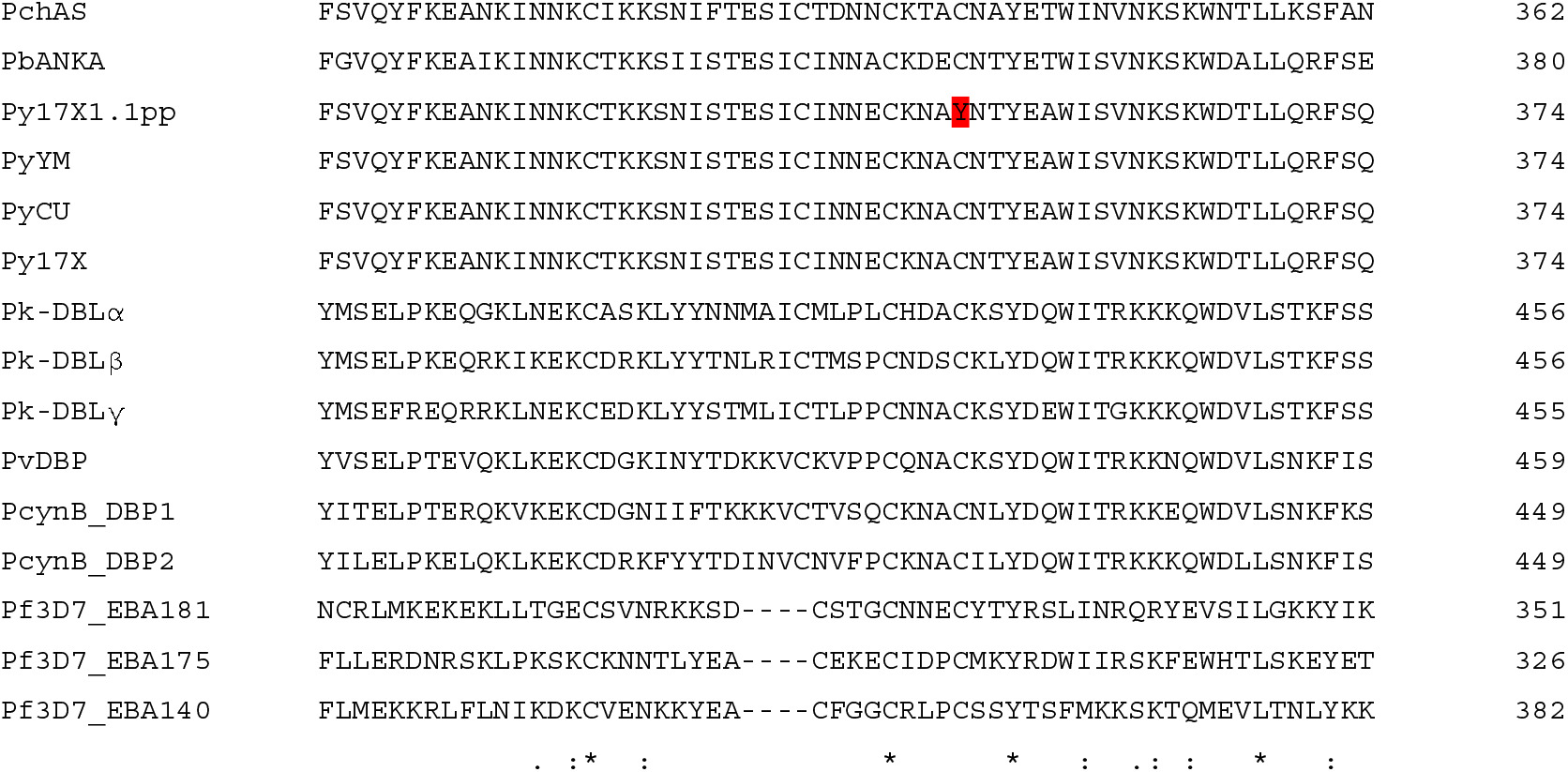
EBL Amino acid sequence alignment of various malaria species and *P. yoelii* strains. EBL orthologous and paralogous sequences from a variety of malaria species and *P. yoelii* strains were aligned using ClustalW. Only the amino acids surrounding position 351 are shown. The cysteine in positon 351 in *P. yoelii* is highly conserved across strains and species, with only strain 17X1.1pp bearing a C→ Y substitution. PchAS: *Plasmodium chabaudi* AS strain; PbANKA: *Plasmodium berghei* ANKA strain; Py17X/17X1.1pp/CU/YM: *P. yoelii* 17X,17X1.1pp,CU,YM strains; Pk-DBL α/β/γ: *Plasmodium knowlesi* Duffy Binding Ligand α/β/γ (H strain); PvDBP: *Plasmodium vivax* Duffy Binding Protein (Sal-I strain); PcynB_DBP1/2: *Plasmodium cynomolgi* Duffy Binding Proteins 1/2 (B strain); Pf3D7_EBA140/175/181: *Plasmodium falciparum* Erythrocyte Binding Antigens 140/175/181 (3D7 strain).

In order to study the role of the mutation, the *Pyebl* gene locus of slow growing CU and faster growing 17X1.1pp clones were replaced with the alternative allele, as well as with the wild-type allele. To establish whether the C351Y substitution affected EBL localization, as was shown for the previously described region 6 mutation, IFA was performed. This revealed that, unlike the mutation in region 6 described previously (23), the EBL proteins of 17X1.1pp and CU were both found to be located in the micronemes (**Figure 5** and **Supplementary Figure S2**).

**Figure 5.**
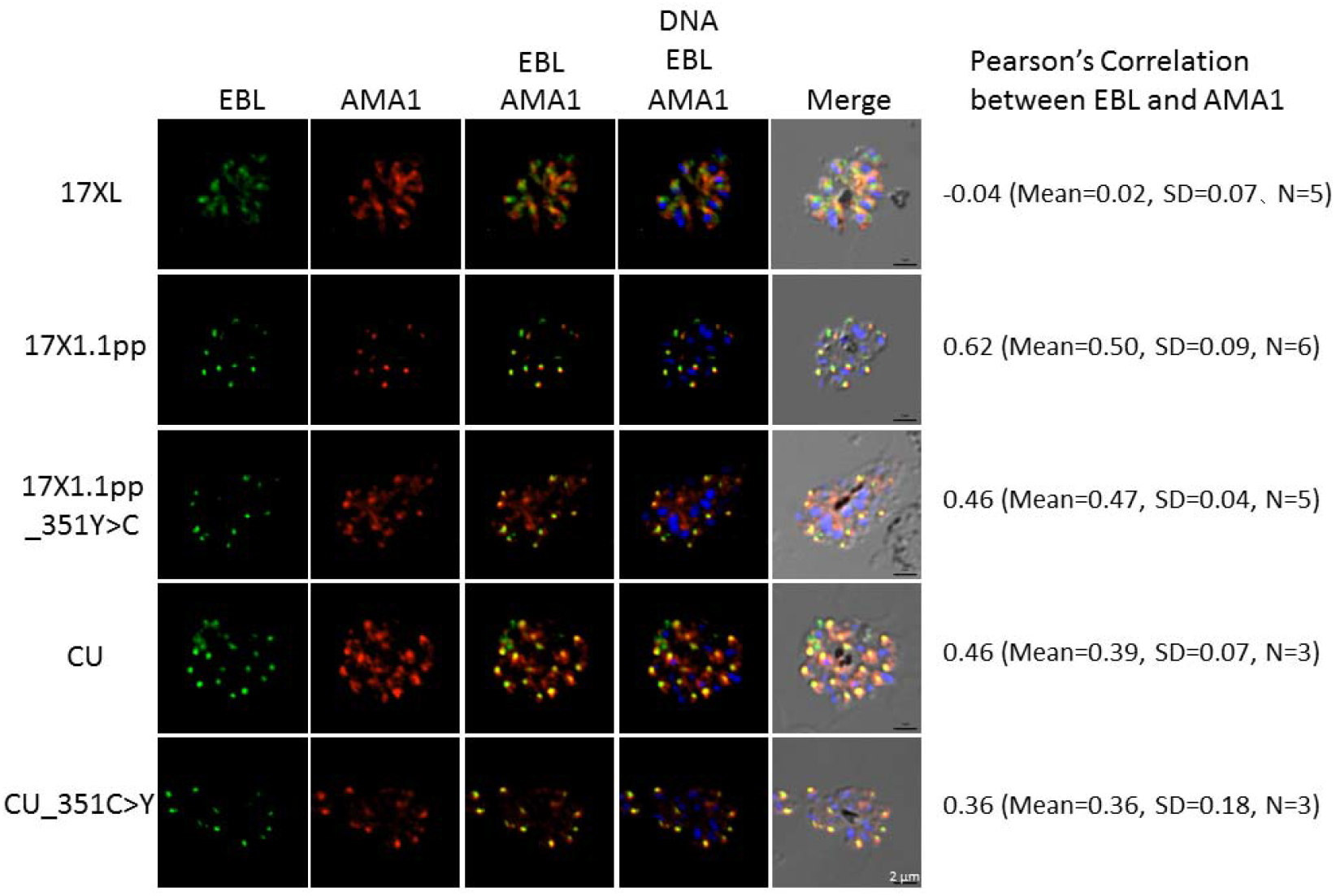
The C351Y mutation does not affect EBL subcellular localization in *P. yoelii. P. yoelii* schizonts of wild type and transgenic parasite lines were incubated with fluorescent mouse anti-PyEBL serum, fluorescent rabbit anti-AMA1 serum, and DAPI nuclear staining. Colors indicate the localization of the PyEBL (green) and AMA-1 (red) proteins, as well as nuclear DNA (blue). 17XL: fast growing 17X clone previously shown to traffic EBL to the dense granules, not the micronemes, 17X1.1pp: 17 × 1.1pp strain, CU: CU strain, 17X1.1-351Y > C: 17X1.1pp strain transfected with the CU allele for PyEBL, CU-351C > Y: CU strain transfected with the 17X1.1pp allele of PyEBL. Pearson’s correlation values on the left of the figure were calculated using between 3 and 6 samples as indicated using the cellular location of the PyEBL (EBL) and AMA-1 proteins. The mean correlation values vary between 0.36 and 0.62 for all samples, except sample 17XL (-0.04). This indicates a shift in the location of PyEBL occurring in 17XL, but not in any of the other parasite lines.

Transgenic clones were grown in mice for 10 days alongside wild-type clones. Pair-wise comparisons between transgenic clones showed that allele substitution could switch growth phenotypes in both strains (**Figure 6A and 6B**). RNA-seq analysis revealed that transfected gene alleles were expressed normally (**Supplementary Figure S3**). This confirmed the role of the C351Y mutation as underlying the observed growth rate difference.

**Figure 6.**
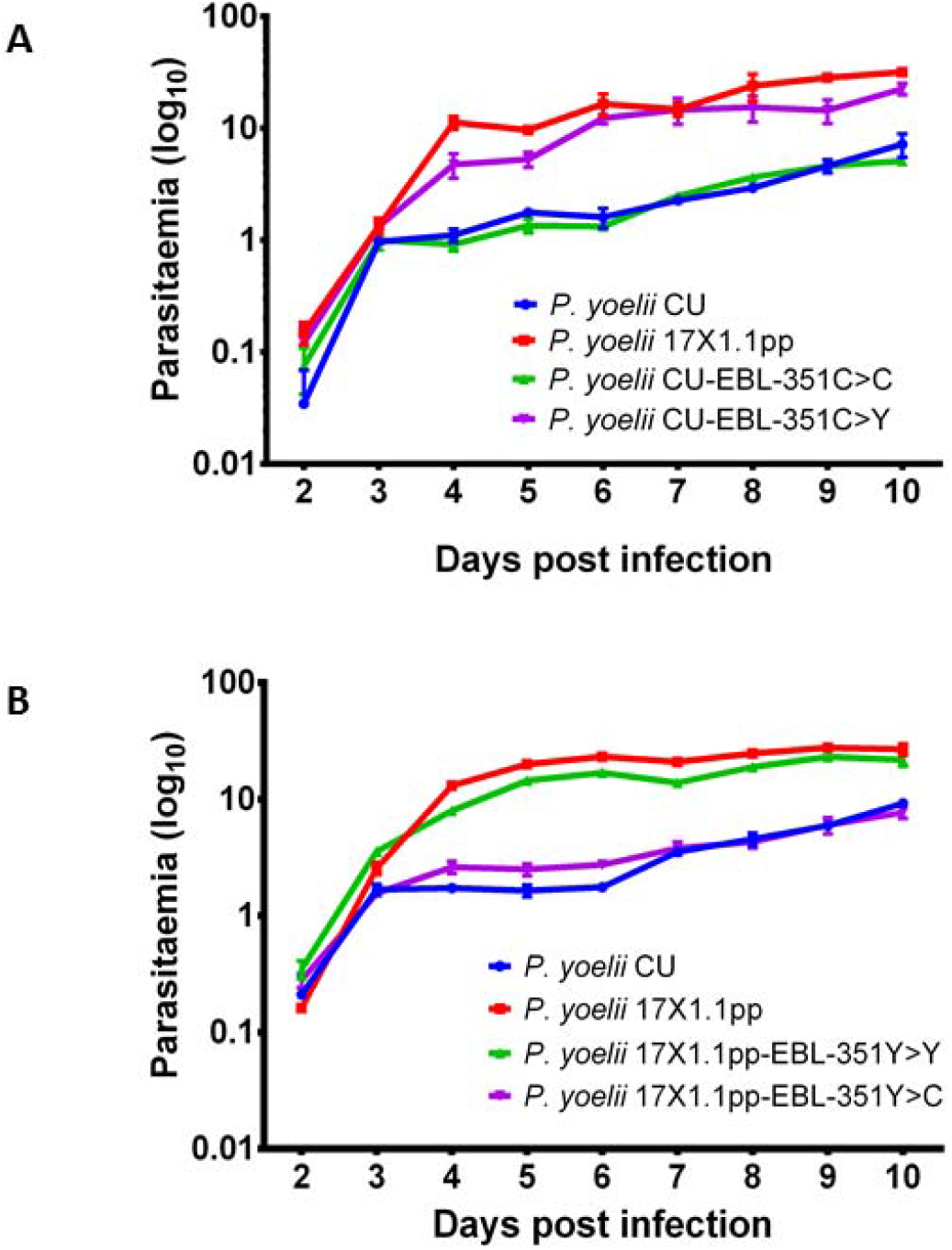
(**A**) Growth rate of *P. yoelii* strains 17X1.1pp, CU and of the CU-strains transfected with either CU (CU-EBL-351C > C) or 17X1.1 (CU-EBL-351C > Y) PyEBL alleles in CBA mice inoculated with 1x10^6^ iRBCs on Day 0. (**B**) Growth rate of *P. yoelii* strains 17X1.1pp, CU and of the 17X1.1pp- strains transfected with either 17X1.1 (17X1.1pp-EBL-351Y > Y) or CU (17X1.1pp-EBL-351Y > C) PyEBL alleles in CBA mice inoculated with 1x10^6^ iRBCs on Day 0. Transfection with the 17X1.1pp (EBL-351Y) allele produces a significantly increased growth rate in the CU strain (CU-EBL-351C > C vs CU-EBL-351C > Y: p < 0.01, Two-way ANOVA with Tukey post-test correction) that is not significantly different from 17X1.1pp growth rate following transfection with its native allele (17X1.1pp-EBL-351Y > Y vs. CU-EBL-351C > Y: p > 0.05, Two-way ANOVA with Tukey post-test correction). Conversely, transfection with the CU (EBA-351C) allele significantly reduces growth (17X1.1pp-EBL-351Y > Y vs 17X1.1pp-EBL-351Y > C: p < 0.01, Two-way ANOVA with Tukey post-test correction) and produces a phenotype that is not significantly different from CU transfected with its own allele (CU EBL-351C > C vs 17X1.1pp-EBL-351Y > C: p > 0.05, Two-way ANOVA with Tukey post-test correction).

## DISCUSSION

The development of LGS has facilitated functional genomic analysis of malaria parasites over the past decade. In particular, it has simplified and accelerated the detection of loci underlying selectable phenotypes such as drug resistance, SSI and growth rate (12, 14, 24). Here we present a radically modified LGS approach that utilizes deep, quantitative WGS of parasite progenies and the respective parental populations, multiple crossing and mathematical modeling to identify selection valleys. This enables the accurate definition of loci under selection and the identification of multiple genes driving selectable phenotypes within a very short space of time. This modified approach allows the simultaneous detection of genes underlying multiple phenotypes, including those with a multigenic basis.

We applied this modified LGS approach to a study of SSI and growth rate in *P. yoelii*, a rodent malaria parasite. We were able to identify three selection valleys that contained three strong candidate genes controlling both phenotypes. Two selection valleys were implicated in SSI; the first time LGS has identified multigenic drivers of phenotypic differences in malaria parasites. Differences in growth rates between the two clones were associated with a single selection valley containing *Pyebl*, which was then shown by the allelic replacement to be the gene controlling this phenotype.

SSI was shown to be controlled by genes within two selection valleys, the strongest on Chr VIII and the other on Chr VII. The Chr VIII valley contained the gene encoding MSP1, a well characterized surface antigen that has been the basis of several vaccine studies ^23^, and which has previously been implicated in SSI in *P. chabaudi* (14, 15). A secondary putative selection valley was detected on Chr VII, which contained several genes with the signature of a potential antigen and that will require further analysis.

The two parental clones CU and 17X1.1pp differ in their growth rates. This is due to the ability of 17X1.1pp to invade both reticulocytes and normocytes, while CU is restricted to reticulocytes (21). A selection valley on Chr XIII was linked to the growth phenotype, and included *Pyebl*, a gene previously implicated in growth rate differences between different strains of *P. yoelii*, due to a mutation in Region 6 of the gene that altered its trafficking, so that the protein was located in the dense granules rather than the micronemes (23, 24). Direct sequencing of the *pyebl* gene of 17X1.1pp and CU revealed a SNP in region 2 of the gene, whereas region 6 displayed no polymorphism. Consistent with this, the EBL proteins of 17X1.1pp and CU were both located in the micronemes, indicating that protein trafficking was unaffected by the region 2 mutation. Allelic replacement of the parasite strains with the alternative allele resulted in a switching of the growth rate to that of the other clone, thus confirming the role of the mutation.

Region 2 of the PyEBL orthologues of *P. falciparum* and *Plasmodium vivax* (25–27) is known to interact with receptors on the red blood cell (RBC) surface. Furthermore the substitution falls within the central portion of the region, which has been previously described as being the principal site of receptor recognition in *P. vivax* (27). Wild-type strains of *P. yoelii* (such as CU) preferentially invade reticulocytes but not mature RBCs, whereas highly virulent strains are known to invade a broader repertoire of RBCs (28). It is thus likely that the C351Y mutation directly affects binding to the host receptor, which in turn may affect RBC invasion preference.

In summary, qSeq-LGS offers a powerful and rapid methodology for identifying genes or non-coding regions controlling important phenotypes in malaria parasites and, potentially, in other apicomplexan parasites. Through bypassing the need to clone and type hundreds of individual progeny, and by harnessing the power of genetics, genomics and mathematical modeling, genes can be linked to phenotypes with high precision in a matter of months, rather than years. Here we have demonstrated the ability of qSeq-LGS to identify multiple genetic drivers underlying two independent phenotypic differences between a pair of malaria parasites; growth rate and SSI. This methodology has the potential power to identify the genetic components controlling a broad range of selectable phenotypes, and can be applied to studies of drug resistance, transmissibility, virulence, host preference, *etc*., in a range of apicomplexan parasites.

## Materials and Methods

### Parasites, mice and mosquitoes

*P. yoelii* CU (slow growth rate) and 17X1.1pp (intermediate growth rate) strains were maintained in CBA mice (SLC Inc., Shizuoka, Japan) housed at 23°C and fed on maintenance diet with 0.05% para-aminobenzoic acid (PABA)-supplemented water to assist with parasite growth. *Anopheles stephensi* mosquitoes were housed in a temperature and humidity controlled insectary at 24°C and 70% humidity, adult flies being maintained on 10% glucose solution supplemented with 0.05% PABA.

### Testing parasite strains for growth rate and SSI

Parasite strains were typed for growth rate in groups of mice following the intravenous inoculation of 1x10^6^ iRBCs of either CU, 17X1.1pp or transfected clones per mouse and measuring parasitaemia over 8-9 days. In order to verify the existence of SSI between the CU and 17X1.1pp strains, groups of five mice were inoculated intravenously with 1x10^6^ iRBCs of either CU or 17X1.1pp parasite strains. After four days, mice were treated with mefloquine (20 mg/kg/per day, orally) for four days to remove infections. Three weeks post immunization, mice were then challenged intravenously with 1x10^6^ iRBCs of a mixed infection of 17X1.1pp and CU parasites. A group of five naive control mice was simultaneously infected with the same material. After four days of growth 10 μl of blood were sampled from each mouse and DNA extracted. Strain proportions were then measured by Real Time Quantitative PCR (RTQ-PCR) using primers designed to amplify the MSP-1 gene (29). All measurements were plotted and standard errors calculated using the Graphpad Prism software (v6.01) (http://www.graphpad.com/scientific-software/prism/). Wilcoxon rank sum tests with continuity correction to measure the SSI effect were done in R (30). Linear mixed model analysis and likelihood ratio tests to test parasite strain differences in growth rate were performed on log-transformed parasitaemia by choosing parasitaemia and strain as fixed factors and mouse nested in strain as a random factor, as described previously (21). Pair-wise comparisons of samples for the transfection experiments were performed using multiple 2-way ANOVA tests and corrected with a Tukey’s post-test in Graphpad Prism.

### Preparations of genetic cross

*P. yoelii* CU and 17X1.1pp parasite clones were initially grown separately in donor mice. These parasite clones were then harvested from the donors, accurately mixed to produce an inoculum in a proportion of 1:1 and inoculated intravenously at 1x10^6^ infected red blood cells (iRBCs) per mouse into a group of CBA mice. Three days after inoculation, the presence of gametocytes of both sexes was confirmed microscopically and mice were anesthetized and placed on a mosquito cage containing ~400 female *A. stephensi* mosquitoes six to eight days post emergence. Mosquitoes were then allowed to feed on the mice without interruption. Seven days after the blood meal, 10 female mosquitoes from this cage were dissected to examine for the presence of oocysts in mosquito midguts. Seventeen days after the initial blood meal, the mosquitoes were dissected, and the salivary glands (containing sporozoites) were removed. The glands were placed in 0.2–0.4 mL volumes of 1:1 foetal bovine serum/Ringer’s solution (2.7 mM potassium chloride, 1.8 mM calcium chloride, 154 mM sodium chloride) and gently disrupted to release sporozoites. The suspensions were injected intravenously into groups of CBA mice in 0.1 mL aliquots to obtain blood stage *P. yoelii* CU 17X1.1pp cross progeny.

### Selection of uncloned cross progeny for linkage group selection analysis

The recombinant progeny from the above cross was subjected to growth rate selection pressure by passage in naïve mice for four days. The resulting population after growth rate selection was referred to as the “growth rate selected cross progeny”. For immune selection, mice immunized with blood stage parasites of either *P. yoelii* CU or 17X1.1pp through exposure and drug cure (as above) were inoculated intravenously with 1x10^6^ parasitized-RBC (pRBC) of the uncloned cross progeny, as described above. The blood stage parasites from the cross progeny grown in these mice which had been previously immunized with CU or 17X1.1pp, were designated “CU-immune selected cross progeny” or “17X1.1pp-immune selected cross progeny”, respectively. The resulting infections were followed by microscopic examination of thin blood smears stained with Giemsa’s solution.

### DNA and RNA isolation

Parental strains and growth rate- or immune-selected recombinant parasites were grown in naïve mice. Parasite-infected blood was passed through a single CF11 cellulose column to deplete host leukocytes, and the genomic DNA (gDNA) was isolated from the saponin-lysed parasite pellet using DNAzol reagent (Invitrogen, Carlsbad, CA, USA) according to the manufacturer’s instructions. For RNA isolation a schizont-enriched fraction was collected on a 50% Nycodenz solution (Sigma Aldrich) and total RNA was then isolated using TRIzol (Invitrogen).

### PCR amplification and sequencing of *ebl* gene

The open reading frame of *P. yoelii ebl* was PCR-amplified from gDNA using KOD Plus Neo DNA polymerase (Toyobo, Japan) with specific primers designed based on the *ebl* sequence in PlasmoDB (PY17X_1337400). *Pyebl* sequences of CU and 17X1.1pp strains were determined by direct sequencing using an ABI PRISM 310 genetic analyzer (Applied Biosystems) from PCR-amplified products. Sequences were aligned using online sequence alignment software Clustal Omega (https://www.ebi.ac.uk/Tools/msa/clustalo/) provided by EMBL-EBI.

### Whole genome re-sequencing and mapping

*P. yoelii* samples were sequenced using paired end Illumina reads (100 bp) (ENA: PRJEB15102). The paired end Illumina data was first quality trimmed using the software Trimmomatic (31). Illumina sequencing adaptors were first removed from the sequences. Following that, trailing bases from both the 5’ and 3’ ends with less than Q20 were trimmed. Lastly, reads found with an average base quality of less than Q20 within a window size of four bases were discarded. Only reads found with surviving pair after the trimming were used for mapping with BWA (32) version 0.6.1 using standard options onto the publicly available genome of *P. yoelii* 17X strain (May 2013 release; ftp://ftp.sanger.ac.uk/pub/pathogens/Plasmodium/yoelii17X/version_2/May_2013/). The SAM alignment files were converted to BAM using the tool Samtools (33). Duplicated reads were marked and removed using the software tool Picard (http://picard.sourceforge.net).

### SNP calling

Genome-wide SNPs were called using the BAM file obtained for the parental CU strain using a previously prepared custom made Python script (18). The script functions as a wrapper for SAMtools mpileup and parses SNPs calls based on mapping quality and Phred base quality scores. In this experiment the values were set at 30 for mapping quality and 20 for base quality. Also, since the *P. yoelii* genome is haploid and the parental strains are clonal, only SNPs where the proportion of any minor alleles was less than 30% was retained, to exclude possible sequencing errors or genuine but uninformative SNPs. The script produces a tab-delimited, human readable table that shows the total number of reads for each of the four possible nucleotides at each SNP. SNPS were called on both parental strains. CU SNPs were then filtered against the 17X1.1pp SNPs to remove any shared SNP calls. The remaining CU SNPs were then used as reference positions to measure the number of reads for each nucleotide in the genetic crosses produced in this study through another Python script (18). This script produced a final table consisting of read counts for each nucleotide of the original CU SNPs in every sample.

### Mathematical Methods for the Identification of loci under Growth Rate and Immune Selection

SNP frequencies were processed to filter potential misalignment events. We note that, during the cross, a set of individual recombinant genomes are generated. Considering the individual genome *g*, we define the function *a_g_(i)* as being equal to 1 if the genome has the CU allele at locus *i*, and equal to 0 if the genome has the 17X1.1pp allele at this locus.

In any subsequent population, the allele frequency *q(i)* at locus *i* can therefore be expressed as

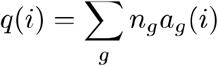

for some set of values *n_g_*, where *n_g_* is the number of copies of genome *g* in the population.

To filter the allele frequencies, we note that each function *a_g_(i)* changes only at recombination points in the genome *g*. As such, the frequency *q(i)* should change relatively slowly with respect to *i* Using an adapted version of code for clonal inference (35), we therefore modelled the reported frequencies *q(i)* as being (beta-binomially distributed) emissions from an underlying diffusion process (denoted by x(i)) along each chromosome, plus uniformly distributed errors, using a hidden Markov model to infer the variance of the diffusion process, the emission parameters, and an error rate. A likelihood ratio test was then applied to identify reported frequencies that were inconsistent with having been emitted from the inferred frequency *x(i)* at locus *i* relative to having been emitted from a distribution fitted using the Mathematica package via Gaussian kernel estimation to the inferred global frequency distribution *{x(i)}*, measured across all loci; this test filters out reported frequencies potentially arising from elsewhere in the genome.

Next, the above logic was extended to filter out clonal growth. In the event that a specific genome g is highly beneficial, this genome may grow rapidly in the population, such that *n_g_* is large. Under such circumstances the allele frequency *q(i)* gains a step-like quality, mirroring the pattern of *a_g_(i)*. Such steps may potentially mimic selection valleys, confounding any analysis.

In order to detect sudden frequency changes arising from clonal growth, a ‘jump-diffusion’ variant of the above hidden Markov model was applied, in which the allele frequency can change either through a diffusion process or via sudden jumps in allele frequency, modelled as random emissions from a uniform distribution on the interval [*0,1*]. For each interval *(i,i + 1)* the probability that a jump in allele frequency had occurred was estimated. Where potential jumps were identified, the allele frequency data was split, such that analyses of the allele frequencies did not span sets of alleles with such jumps. The resulting segments of genome were then analyzed under the assumption that they were free of allele frequency change due to clonal behavior.

Inference of the presence of selected alleles was performed using a series of methods. In the absence of selection in a chromosome, the allele frequency is likely to remain relatively constant across each chromosome. Firstly, therefore, a ‘non-neutrality’ likelihood ratio test was applied to each contiguous section of genome, calculating the likelihood difference between a model of constant frequency *x(i)* and the variable frequency function *x(i)* inferred using the jump-diffusion model. Secondly, an inference was made of the position of the allele potentially under selection in each region. Under the assumptions that selection acts for an allele at locus *i*, and that the rate of recombination is constant within a region of the genome, previous work on the evolution of cross populations (20, 35) can be extended to show that the allele frequencies within that region of the genome at the time of sequencing are given by

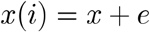

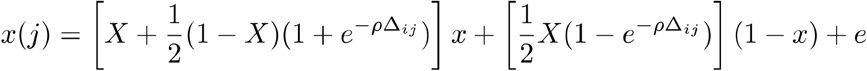

for each locus j not equal to i, where X is the CU allele frequency at the time of the cross,is the local recombination rate, *ρ_ij_* is the distance between the loci *i* and *j*, *x* is an allele frequency, and *e* describes the effect of selection acting upon alleles in other regions of the genome. A likelihood-based inference was used to identify the locus at which selection was most likely to act. In regions for which the ‘non-neutrality’ test produced a positive result for data from both replica crosses, and for which both the inferred locus under selection, and the direction of selection acting at that locus were consistent between replicas, an inference of selection was made.

For regions in which an inference of selection was made, an extended version of the above model was applied, in which the assumption of locally constant recombination rate was relaxed. Successive models, including an increasing number of step-wise changes in the recombination rate, were applied, using the Bayesian Information Criterion (36) for model selection. A model of selection at two loci within a region of the genome was also examined.

Given an inference of selection, a likelihood-based model was used to derive confidence intervals for the position of the locus under selection. A combined confidence interval utilized data from both replicates, representing the extent of uncertainty inherent in the model, on the basis of the modelling assumptions. A second, more conservative, interval was calculated as the union of the two confidence intervals calculated when the model was applied separately to data from each replica; this second approach aims to account for ‘biological noise’, encompassing any deviation in the experimental system from the model assumptions. Full details of the mathematical methods are contained within Supporting Information (**Supplementary Protocol 1**).

### Gene information and NS/S SNP ratios for *P. falciparum* orthologues

For each combined conservative interval of relevant selective sweeps, genes were listed based on the 6.2 PlasmoDB annotation and verified against the current annotation (v26). For each gene, transmembrane domains and signal peptides predicted in PlasmoDB were also indicated. *P. falciparum* orthologues were also indicated, together with their NS/S SNP ratios stored in the current version of PlasmoDB.

### Plasmid construction to modify *P. y. yoelii ebl* gene locus

All primer sequences are given in **Table S7**. Plasmids were constructed using MultiSite Gateway cloning system (Invitrogen). attB-flanked *ebl* gene products, attB12-*Py*CU-EBL.ORF and attB12-*Py*17X1.1pp-EBL.ORF, were generated by PCR-amplifying both *P. yoelii* CU and *P. yoelii* 17X1.1pp *ebl* gene with yEBL-ORF.B1F and yEBL-ORF.B2R primers. attB-flanked *ebl*-3U (attB41-*Py*CU-EBL-3U and attB41-*Py*17X1.1pp-EBL-3U) was similarly generated by PCR-amplifying *P. yoelii* gDNA with yEBL-3U.B4F and yEBL-3U.B1R primers. attB12-*Py*CU-EBL.ORF and attB12-*Py*17X1.1pp-EBL.ORF were then subjected to a separate BP recombination with pDONR™221 (Invitrogen) to yield entry plasmids, pENT12-*Py*CU-EBL.ORF and pENT12-*Py*17X1.1pp-EBL.ORF, respectively. attB41-*Py*CU-EBL-3U and attB41-*Py*17X1.1pp-EBL-3U fragments were also subjected to independent BP recombination with pDONR™P4-P1R (Invitrogen) to generate pENT41-*Py*CU-EBL-3U and pENT41-*Py*17X1.1pp-EBL-3U, respectively. All BP reactions were performed using the BP Clonase™ II enzyme mix (Invitrogen) according to the manufacturer’s instructions. To change *P. yoelii* CU *ebl* gene nucleotide 1052G to 1052A (351Cys to 351Tyr), pENT12-*Py*CU-EBL.ORF entry clone was modified using KOD-Plus-Mutagenesis Kit (TOYOBO) with primers P1.F and P1.R to yield pENT12-*Py*CU-EBL.ORF-C351Y. pENT12-*Py*17X1.1pp- EBL.ORF was also modified from 1052A to 1052G (351Tyr to 351Cys) using primers P2.F and P1.R to yield pENT12-*Py*17X1.1pp-EBL.ORF-Y351C. pHDEF1-mh that contains a pyrimethamine resistant gene selection cassette (37) (a gift from Hernando del Portillo) was digested with *Sma*I and *Apa*I to remove PfHRP2 3′ UTR DNA fragment, cohesive end was blunted, and a DNA fragment containing ccdB-R43 cassette and *P. berghei* DHFR-TS 3′ UTR that was amplified from pCHD43(II) (38) with primers M13R.F3F and PbDT3U.F3R was ligated to generate pDST43-HDEF-F3. pENT12-*Py*CU-EBL.ORF-C351Y and pENT12-*Py*17X1.1pp-EBL.ORF-Y351C entry plasmids were each separately subjected to LR recombination reaction (Invitrogen) with a destination vector pDST43-HDEF-F3, pENT41-*Py*CU-EBL-3U or pENT41-*Py*17X1.1pp-EBL-3U and a linker pENT23-3Ty1 vector to yield replacement constructs pREP-*Py*CU-EBL-C351Y and pREP-*Py*17X1.1pp-EBL-Y351C, respectively. Control constructs pREP-*Py*CU-EBL-C351C and pREP-*Py*17X1.1pp-EBL-Y351Y were also prepared in a similar manner. These LR reactions were performed using the LR Clonase™ II Plus enzyme mix (Invitrogen) according to the manufacturer’s instructions.

### Allele replacement

*P. yoelii* schizont-enriched fraction was collected by differential centrifugation on 50% Nycodenz in incomplete RPMI1640 medium, and 20 µg of *Apa*I- and *Stu*I-double digested linearized transfection constructs were electroporated to ~1x10^7^ of enriched schizonts using a Nucleofector device (Amaxa) with human T-cell solution under program U-33 (39). Transfected parasites were intravenously injected into 7-week-old ICR female mice, which were treated by administering pyrimethamine in the drinking water (conc) 24 hours later for a period of 4-7 days. Before inoculation of *P. yoelii* 17X1.1pp parasite, mice were treated with phenylhydrazine to increase the reticulocyte population in the blood. Drug resistant parasites were cloned by limiting dilution. Integration of the transfection constructs was confirmed by PCR amplification with a unique set of primers (see table S10) for the modified *ebl* gene locus, followed by sequencing (both Sanger and NGS). pREP-*Py*CU-EBL-C351Y and pREP-*Py*CU-EBL-C351C were transfected to *P. yoelii* CU parasites whereas pREP-*Py*17X1.1pp-EBL-Y351C and pREP-*Py*17X1.1pp-EBL-Y351Y were transfected to *P. yoelii* 17X1.1pp parasites.

### Phenotype analysis

To assess the course of infection of wild type and transgenic parasite lines, 1x10^6^ pRBCs were injected intravenously into 8-week old female CBA mice. Thin blood smears were made daily, stained with Giemsa’s solution, and parasitaemias were examined microscopically.

### RNA-Seq

Whole blood from mice infected with *P. yoelii* on day 5 post-infection were host WBC depleted and saponin lysed to obtain the parasite pellet. Total RNA was extracted using TRIzol reagent. Strand-specific RNA sequencing was performed from total RNA using TruSeq Stranded mRNA Sample Prep Kit LT according to manufacturer’s instructions. Libraries were sequenced on an Illumina HiSeq 2000 with paired-end 100 bp read chemistry (ENA: PRJEB15102). RNA-seq reads were mapped onto Plasmodium yoeli 17X version 2 from GeneDB (http://www.genedb.org) using TopHat 2.0.13 (40) and visualized using Artemis genome visualization tool (41).

### Indirect immunofluorescence assay

Schizont-rich whole blood was obtained from *P. yoelii* infected mouse tail and prepared air-dried thin smears on glass slides. The smears were fixed in 4% paraformaldehyde containing 0.0075% glutaraldehyde (Nacalai Tesque) in PBS at room temperature (RT) for 15 min, rinsed with 50 mM glycine (Wako) in PBS. Samples were permeabilized with 0.1% Triton X–100 (Calbiochem) in PBS for 10 min, then blocked with 3% BSA (Sigma) in PBS at RT for 30 min. Next, samples were immunostained with primary antibodies using mouse anti–*Py*EBL (23) at 37°C for 1 h. This was followed by 3 washes with PBS then incubation with Alexa Fluor^®^ 488 goat anti–mouse and Alexa Fluor® 594 goat anti–rabbit antibodies (Invitrogen; final 1:1000) in 3% BSA in PBS at 37°C for 30 min. Parasite nuclei were stained with 4, 6-diamidino-2-phenylindole (DAPI; Invitrogen, final 0.2 µg/mL). Stained parasites were mounted with Prolong Gold® antifade reagent (Invitrogen). Slides were visualized using a fluorescence microscope (Axio imager Z2; Carl Zeiss) with 100x oil objective lens (NA 1.4, Carl Zeiss). Images were captured using a CCD camera (AxioCam MRm; Carl Zeiss) and imaged using AxioVision software (Carl Zeiss). Pearson’s colocalization coefficients were calculated using Zen software (Carl Zeiss).

## Acknowledgements

This work was supported by grants from the Naito Foundation (to R.Cu); the JSPS (project numbers Nos. JP25870525, JP24255009 and JP16K21233) (to R.Cu), A Royal Society Bilateral Grant for Co-operative Research (to R.Ca and R.Cu) and a Sasakawa Foundation Butterfield Award (to R.Cu), faculty baseline funding from the King Abdullah University of Science and Technology (KAUST) to AP, and Grants-in-Aids for Scientific Research on Innovative Areas JR23117008, MEXT, Japan (to OK). CJRI was supported by a Sir Henry Dale Fellowship, jointly funded by the Wellcome Trust and the Royal Society (Grant Number 101239/Z/13/Z). This work was conducted in part at the Joint Usage / Research Center on Tropical Disease, Institute of Tropical Medicine, Nagasaki University. We thank Ho Y. Shwen for initial contributions to the project and Andrej Fisher for discussions and for the provision of code used in the jump-diffusion analysis. AM is supported by GI-CoRE funded to the Research Center for Zoonosis Control in Hokkaido University.

